# Beyond APs and VPs: evidences of chaos and self-organized criticality in electrical signals in plants

**DOI:** 10.1101/064790

**Authors:** G. F. R. Saraiva, A. S. Ferreira, G. M. Souza

## Abstract

Studies on plant electrophysiology are mostly focused on specific traits of action potentials (APs) and/or variation potentials (VPs), often in single cells. Inspired by the complexity of the signaling network in plants and by analogies with some traits of neurons in human brains, we have sought for evidences of high complexity in the electrical dynamics of plant signaling, beyond APs and VPs responses. Thus, from EEG-like data analyses of soybean plants, we showed consistent evidences of chaotic dynamics in the electrical time series. Furthermore, we have found that the dynamic complexity of electrical signals is affected by the plant physiological conditions, decreasing when plant was stressed. Surprisingly, but not unlikely, we have observed that, after stimuli, electrical spikes arise following a power law distribution, which is indicative of self-organized criticality (SOC). Since, as far as we know, these were the first evidences of chaos and SOC in plant electrophysiology, we have asked follow-up questions and we have proposed new hypotheses, seeking for improving our understanding about these findings.

## Introduction

Because of the sessile and modular nature and the continuing challenges on their surviving, the plants must be able to perceive, interpret and respond to various environmental stimuli by integrating the signals received by many different sensors in the cells. To this end, they use a system that involves a complex network of signal transduction, involving cell-cell and long distance communication, enabling integration of their body parts (modules) as a whole, providing the ability to adjust their phenotype to different environmental conditions^1,2,3^.

Different types of signals, such as hormones, ROS, Ca^2+^ and electrical signals, compose the plant’s signaling network^3,4^. Specifically, there are three types of electrical signals in plants: APs (action potentials), VPs (variation potentials), and SPs (system potentials). Strong evidences have demonstrated that these signals play central role in both cell-cell and long-distance communication in plants^5–8^.

APs are characterized by spike-like changes of the resting membrane potential and, independent of the stimulus strength, starts propagating through the plant with a defined amplitude and velocity. Like in animals, APs seem to be all-or-nothing events^9,10^. VPs differ from APs in various ways. VPs do not obey the all-or-nothing law, they are known as slow wave potentials (SWPs) with variable shape, amplitude and time frame. Moreover, the signals are related with the stimulus strength, and last for periods of 10 s up to 30 min^11,12^. System potentials (SPs), in contrast to APs and VPs, reflect a systemic self-propagating hyperpolarization of the plasma membrane or depolarization of the apoplastic voltage. Like VPs, SPs have a magnitude and duration that are depended on the stimulus, but they are initiated via membrane hyperpolarization through the sustained activation of the proton pump. SPs are dependent of experimental conditions, and then they may occur under a very specific set of environmental conditions^4,11^.

Despite of different induction mechanisms, the electrical signals are able to inform distant cells about local stimuli, triggering proper physiological responses to a multitude of environmental stimuli^6^. As far as we know, the studies linking electrical signaling with physiological responses to environmental cues are based on the analysis of APs or VPs waves, taking into account only parameters as frequency, amplitude, distance of propagation and time frame^6,8,12,13^.

However, quite often mixed electrical potential waves (EPWs) are recorded, for instance, as result of overlapping APs and VPs, which impedes proper signal analysis^14^. It is also important to note that even though APs, VPs and SPs most likely occur through distinct molecular mechanisms, these phenomena can arise within similar temporal scales, creating a complex web of systemic information in which several electrical signals may be layered on top of each other in time and space^4^. Accordingly, Masi et al. (ref 15) have reported strong evidence of synchronization of electrical spikes among different cells in the maize root apex, suggesting a collective behavior of groups of cells interconnected by the plasmodemata network. Therefore, the temporal and spatial dynamics of electrical signaling in plants exhibit complex behavior. Indeed, Cabral et al. (ref 16) showed evidence of high complexity dynamics in plant electrical signals, exhibiting a large spectrum of frequencies, but not random.

Thus, we have hypothesized that the complexity of the massive ionic flow through plant tissues could be similar to the complexity observed in the human brain, since they share analogous (but non homologous) mechanisms of membrane depolarization/re-polarization based on transmembrane ion fluxes, linking electrically a plenty of cells and engendering a complex network of information transmission^4,17^.

Actually, since the mid 1980s, scientists began to apply chaos theory to human electroencephalogram (EEG) data sets, uncovering non-random complex patterns underlying brain electrical signals. Accordingly, EEG signals can be interpreted as the output of a deterministic system of relatively low complexity, but containing highly non-linear elements^18^. Non-linear dynamical systems can exhibit chaotic behavior and, thus, become unpredictable over a long time scale^19^.

In animals, particularly humans, studies on the temporal dynamics of electric signals obtained by EEG are allowed to establish consistent relationship between the state of health of individuals and the complexity measures in EEG, applying nonlinear time series analyses techniques based on deterministic chaos theory^18,20–22^. Relationships between the intricate variation in the physiological time-series parameters and the state of plants under different environmental conditions were previously studied by Souza et al. (refs 23-25) allowing to hypothesize that eventual changes in the dynamics of EEG-like signals in plants could be associated to environmental stimuli responses.

Gardiner^26,27^ has suggested the possibility of using EEG in plants to relate likely changes in temporal dynamics with different physiological states. But as far as we know, there are no studies in this direction. Thus, the main objective of this study was to investigate the eventual existence of complex information underlying plant electrical signaling, beyond APs and VPs, by analyzing the electrical signals measured with EEG-like technique. Moreover, we have taken in consideration the possibility that stressful environmental stimuli induce changes in the signals complexity.

## Material and Methods

### The plant model and growth conditions

In this study, soybean [*Glycine max* (L) Merrill] cv. Intact Bt/RR was used as plant model. *G. max* offers a good experimental model because it is easily cultivated at laboratory conditions, the available knowledge of its physiology under different environmental stressful conditions and it has been used previously in bioelectrical studies^8^.

Seedlings were obtained from seeds germinated in 180 mL pots with vermiculite as substrate. The pots were kept in maximum capacity of water retention during both germination and early seedling development under controlled conditions (Phytotron EL-101, Eletrolab, Brazil): day/night temperature of 28/22 °C, respectively, 14h of photoperiod with 500 μmol photons m^-2^ s^-1^, and air humid around 60%. The plants were irrigated daily with ½ strength Hoagland nutrient solution, preventing, at same time, both starvation and salinization. The amount of irrigation was determined after weighing the pots with its maximum water retention and verifying the daily evaporation loss.

### Data acquisition and experimental design

For acclimation, one the day before each experimental session, sub-dermal needle electrodes (model EL452, Biopac Systems, US) were inserted into the region between the stem and the roots, below the first pair of simple leaves, using plants with 15 days after germination. Acclimation to the electrodes is required because insertion induces action potentials and local fluctuations in potential variation, which is stabilized in a few hours with the disappearance of action potentials^28^. Each pair of electrodes (positive/negative) was inserted into the stem at a fixed distance (1 cm from each other) and 2-3 mm in depth, ensuring contact with the conducting vessels of the plants. The parts of the electrodes outside the plants were isolated from each other by a block of polystyrene. A third electrode was attached to the structure of the Faraday cage in order to obtain adequate electrical grounding. In each experimental session, data were collected from four plants simultaneously, using a total of forty plants.

All bioelectric measurements were taken in a Faraday cage properly grounded to prevent electric field from the laboratory environment. The bioelectric time series were recorded using a device of electronic acquisition with four channels (model MP36, Biopac Systems, US) with high input impedance. The sampling rate used was 125 Hz with two filters, one high-pass (0.5 Hz cutoff frequency) and a low-pass (1.5 kHz cutoff frequency). The signals were amplified with a gain of 20,000 x, allowing high resolution to capture the voltage variations before and after stimuli, in order to perform suitable and reliable non-linear time series analyses. Data gathering was carried out continuously during 1h before and 1h after application of the environmental stimuli.

The plants were osmotically stimulated with a mannitol solution with water potential of -2.0 MPa, applied directly on the substrate of the pots, taking care to avoid any mechanical contact with the plant. According to several previous studies, this osmotic potential is stressful, but no lethal, for soybean plants^29,30^. The temperature of the mannitol solution was previously kept in equilibrium with the temperature inside the Phytotron to avoid additional disturbances on the root system.

### Mathematical data analyses

In this study, we analyzed the time series *ΔV*={Δ*v*_1_, *Δv*_2_,…,*Δv*_*N*_}, where Δ*v_i_* is the potential difference between the electrodes inserted in the plants, as described above. The series analyzed correspond to samplings with the total length N ∼75,000 points, corresponding to 600 s of the data measured before and after stimuli.

However, before experimental data analyzes, the noise from the EEG device was characterized as follows. First, it was calculated the Pearson linear correlation function *ρ(τ)* between two random variables *x* and *y*, delayed *τ* each other, by the equation

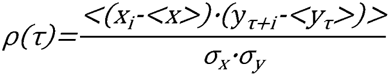

where -1 ≤ *ρ(τ)* ≤ 1, τ is the time lag and *<…>* represents an average; 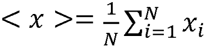, 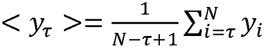, 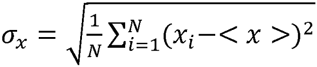; and 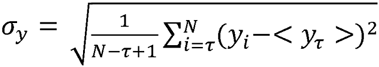. For *ρ(τ)>0* the variables are positively correlated, and for *ρ(τ)<0* the variables are negatively correlated throughout time. For *ρ(τ)=0* the variables are uncorrelated. When *ρ(τ)* decay exponentially the correlation have short range, showing well defined characteristic time correlation. On the other hand, when *ρ(τ)* follows a power law, the correlations have long range without characteristic time. Considering *x*=*Δv* and *y*=*Δv* in *(ρ(τ))*, we have the auto-correlation function for the variable Δ*v* with lag *τ*.

In the *ρ(τ)* calculated for data from de open electrode (noise of the device) it was observed a exponential decay 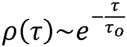, meaning that the correlations were short range with characteristic time τ_o_= 0.08s. The probability density function (*pdf*) of the noise was Gaussian, as indicated by the *normal probability plot* method^31^ (Figure 1, inset). This method consists of linearize the Gaussian of the *pdf*. When the data analyzed are arranged in the line, implies that they follow the same distribution. The Gaussian distribution of the noise was corroborated by the analysis of skewness (β) and kurtosis (κ). Gaussian *pdf* shows symmetrical (β = 0.0) and peaked distribution with κ = 3.0. Hence, we have demonstrated that the EEG device has a Gaussian noise (Figure 1), do not interfering significantly with the quality of the dynamics of the signals measured.

**Figure 1.**
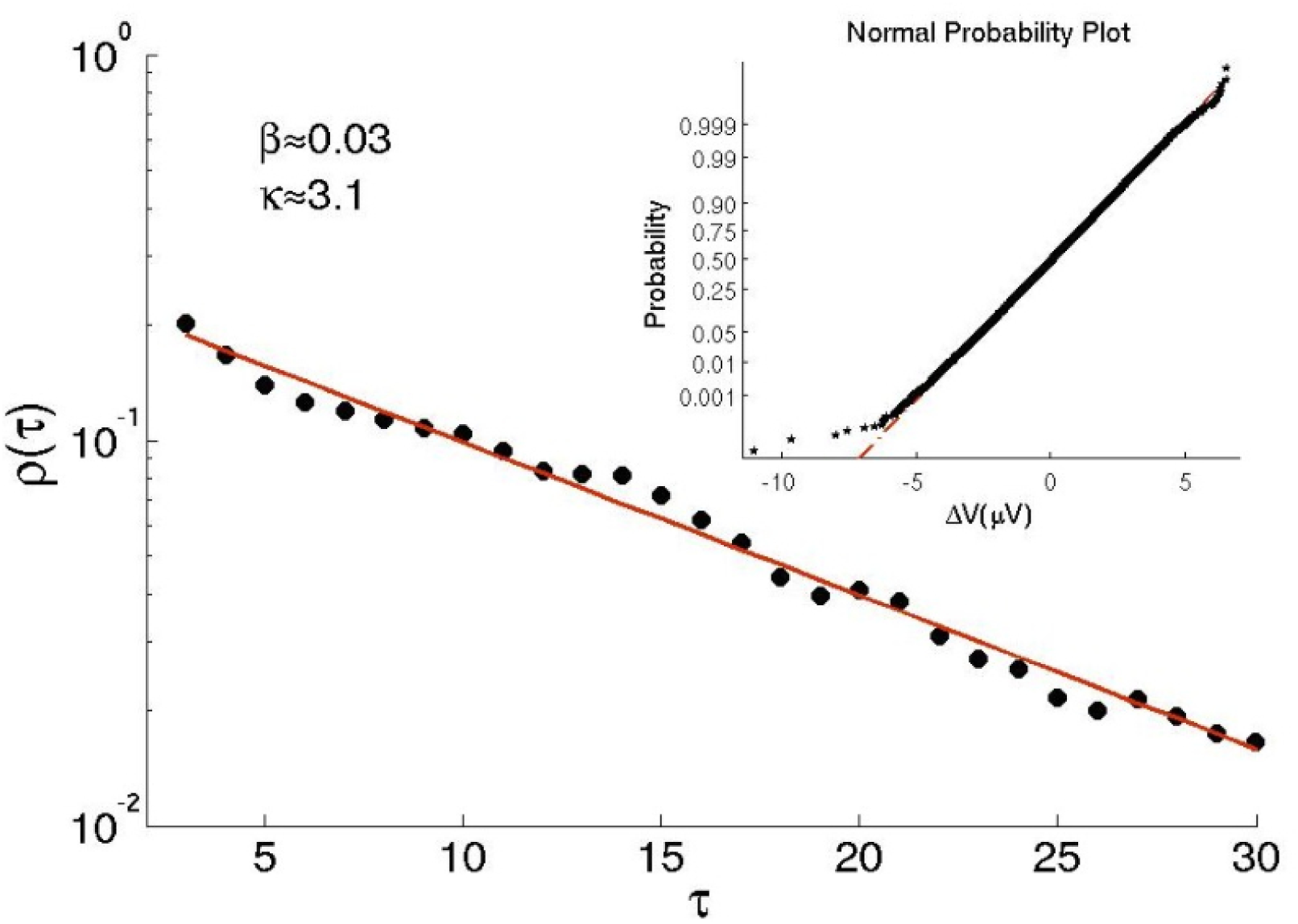
The autocorrelation function *ρ(τ)*with exponential decay, indicating that the temporal correlations are short range. The filled symbols are the experimental data and the continuous line is the fitting. In the inset, the normal probability plot test indicates that the *pdf* of the noise of the device is Gaussian. β = skewness; κ = kurtosis.

The time series sampled were analyzed by different methods in order to characterize the temporal dynamic of the signals, focusing in comparing the dynamics before and after the stimuli. First, we analyzed the auto-correlation function for each time series and, then, the cross-correlation function was calculate between the before and after the stimuli series. Second, the complexity of the time series were estimated by computing the Approximate Entropy (*ApEn*) and, then, tested for chaotic behavior by computing the largest Lyapunov exponents of the series.

### Experimental time series analyses

The auto-correlation functions of the time series sampled before and after stimuli were calculate as described above.

The calculation of Approximate Entropy, *ApEn (m,r)*, follows the equation^33^

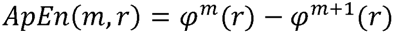

where 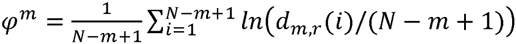 and *d_m,r_*(*i*) is the number (*i*) of vectors pairs m-dimensional to an Euclidian distance less or equal to *r*, and *N-m+1* is the total number of vectors in the embedding dimension *m*. For this study, according to the demonstrations of Pincus^32,33^, we assume *r= 0.2σ* (20% of the standard deviation of the time series ΔV) and *m* = 2.

ApEn has been used as a robust method to measure the complexity (irregularity level) of biological time series. ApEn assigns a non-negative number to a sequence or time-series, with larger values corresponding to greater apparent process randomness or serial irregularity, and smaller values corresponding to more instances of recognizable features or patterns in the data^25,32,33^.

### Largest Lyapunov exponent

To characterize chaos in the time series, the largest Lyapunov exponent was calculated according to Rosenstein’s method^34^. The Lyapunov exponent^35^ quantifies the rate of separation of infinitesimally close trajectories in the phase space. Taking *ΔS_o_>0* as the initial distance between two trajectories apart each other by a slight disturbance, the temporal evolution of this separation is given by the equation Δ*S*(*τ*) = Δ*S_o_* · *e^λ·τ^*, where λ is the Lyapunov exponent and τ is the time step. When λ<0 the two trajectories converge and the system is stable. When *λ>0* the trajectories diverge in the space-phase, and the system exhibit a chaotic behavior. Stochastic dynamics, mathematically, show λ→∞ that, in the plot of *log(ΔS(τ))* versus τ, is indicated by a vertical straight line trending in the initial part of the plot (such as in Figure 5, in Results).

## Results

The auto-correlation analysis showed that, contrary the noise of the EEG device, all the experimental time series sampled have a long-range correlation. However, the pattern decay of the *ρ(τ)* of the signal measured before and after the stimuli exhibited different behaviors. While the trend of decay of *ρ(τ)* before stimuli was continuous, the trend after stimuli showed a consistent oscillatory behavior. In order to corroborate the observed differences in the *ρ(τ)* between the signals before and after stimuli, a cross-correlation analysis was carried out, showing no correlation between them (Figure 2).

**Figure 2.**
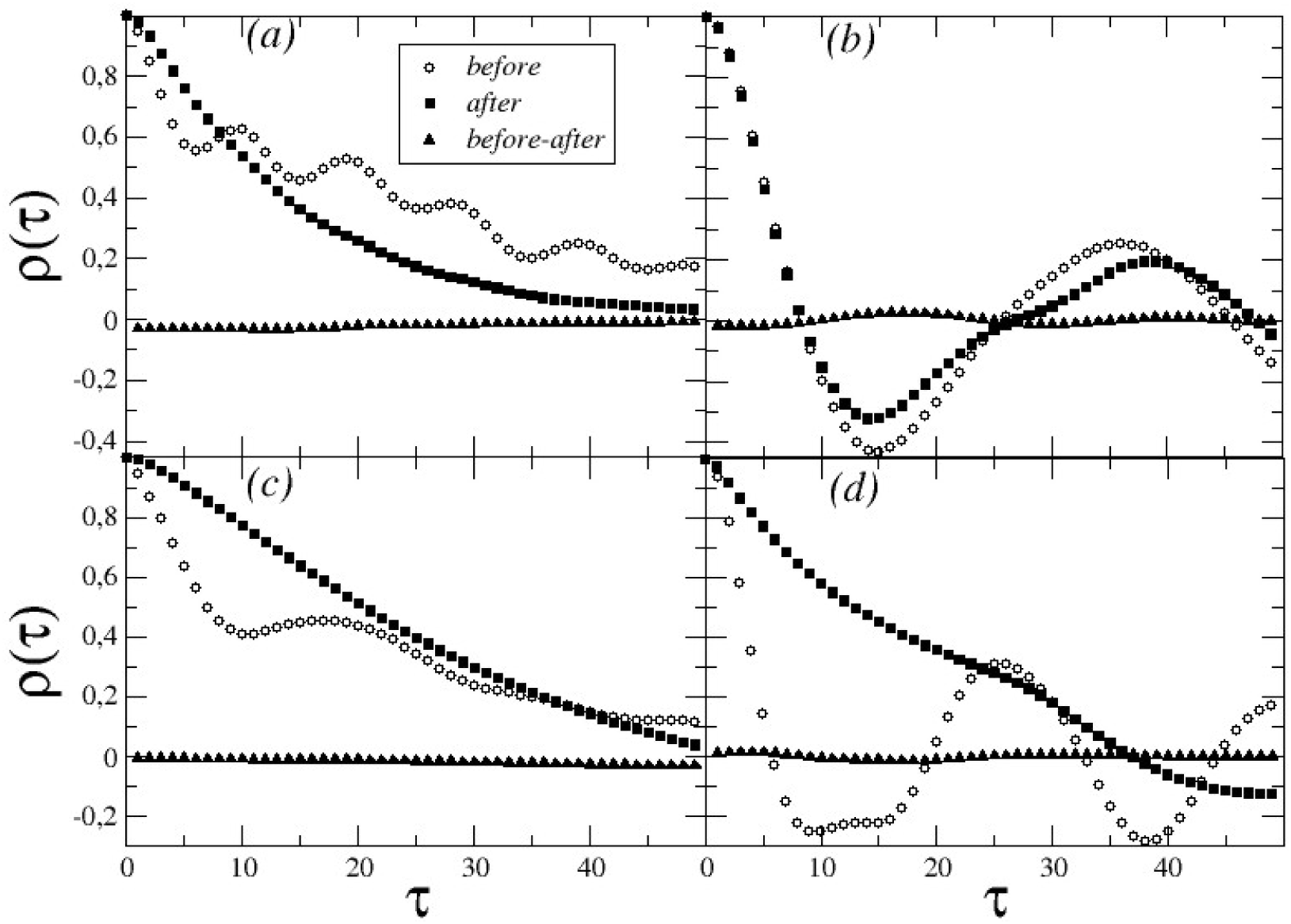
Auto-correlation function *ρ(τ)* of four time series samples (a, b, c, and d) randomly chosen from the total data set (n = 40). The auto-correlation function before and after treatment are represented by empty circles and filled squares, respectively. The cross correlation function before and after stimuli is shown by filled triangles.

The complexity measurements showed consistently that after stimuli there was a decrease in ApEn values (Figure 3), indicating higher complexity in the electrical signals before stimuli. For the most of the series analyzed (after and before stimuli) the Lyapunov exponents were finites and positives, indicating chaotic behavior. Moreover, the Lyapunov exponents were lower after than before stimuli, supporting the ApEn results that have indicated lower complexity in the electrical dynamic after stimuli (Figure 4).

**Figure 3.**
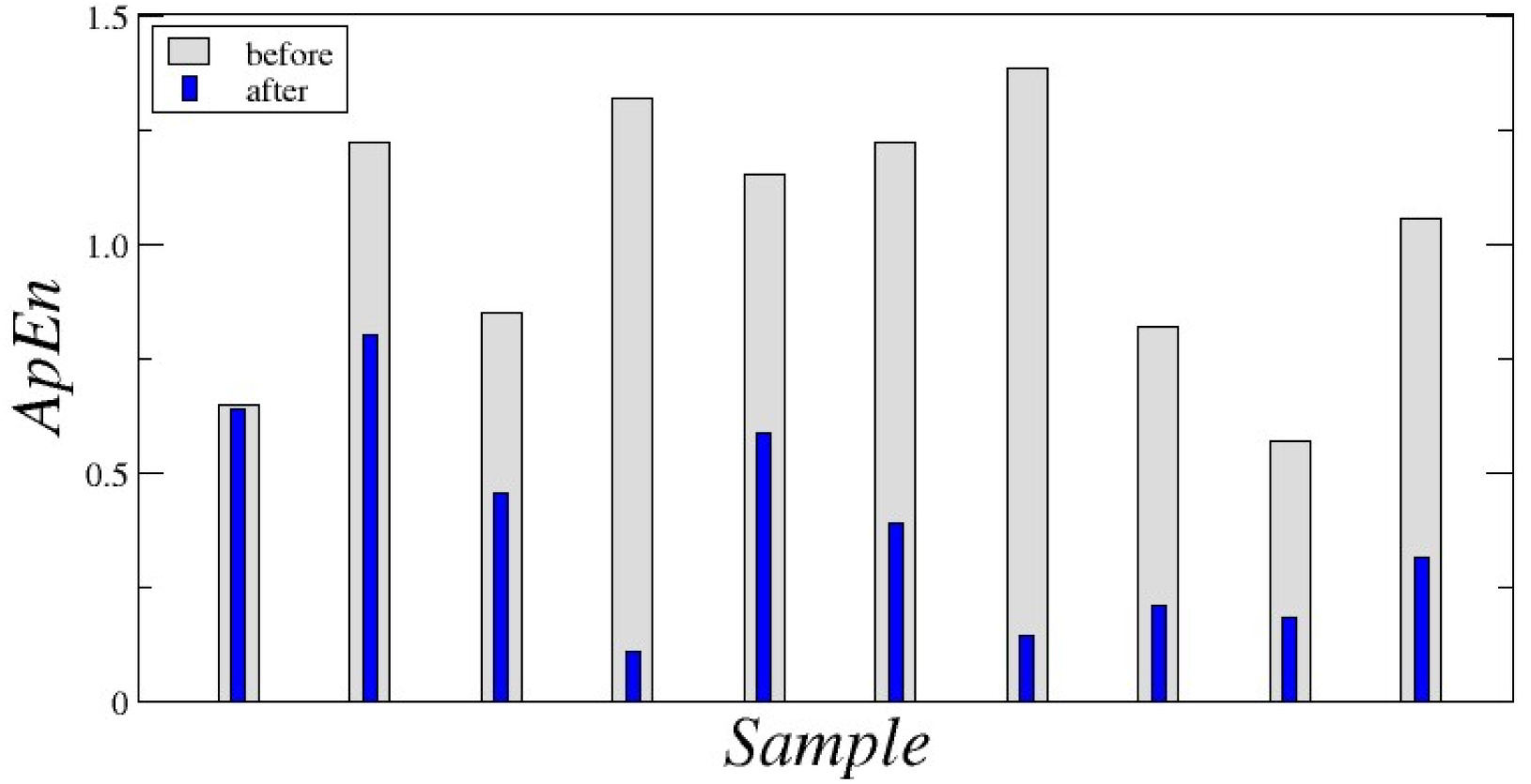
Approximate Entropy (ApEn) before (cross-hatched column) and after (filled column) the stimulus for ten time series samples chosen randomly from the total data set (n = 40).

**Figure 4.**
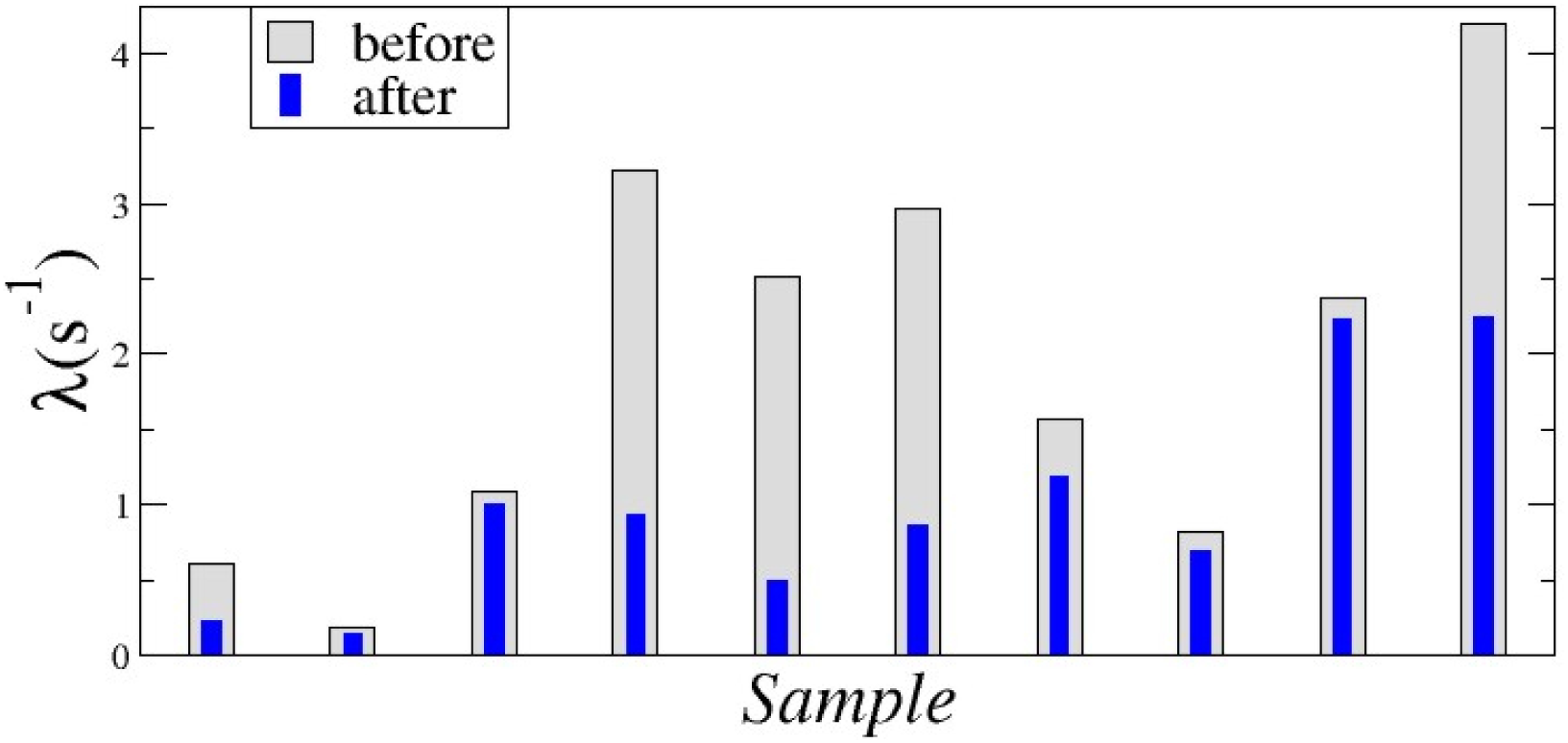
Largest Lyapunov exponent before (cross-hatched column) and after (filled column) of the stimuli for ten time series samples chosen randomly from the total data set (n = 40).

The largest Lyapunov exponent is obtained from the slope of the log of the divergence between two trajectories (*log(Δs) vs Δt*) according to the time step^34^. In some situations observed herein, signals before stimuli showed an asymptotically infinite exponent, suggesting that the series was purely stochastic (Figure 5a). However, in the same plant sampled after the stimulus, the Lyapunov exponent has become finite and positive, indicating that the series has assumed a chaotic dynamics (Figure 5b).

**Figure 5.**
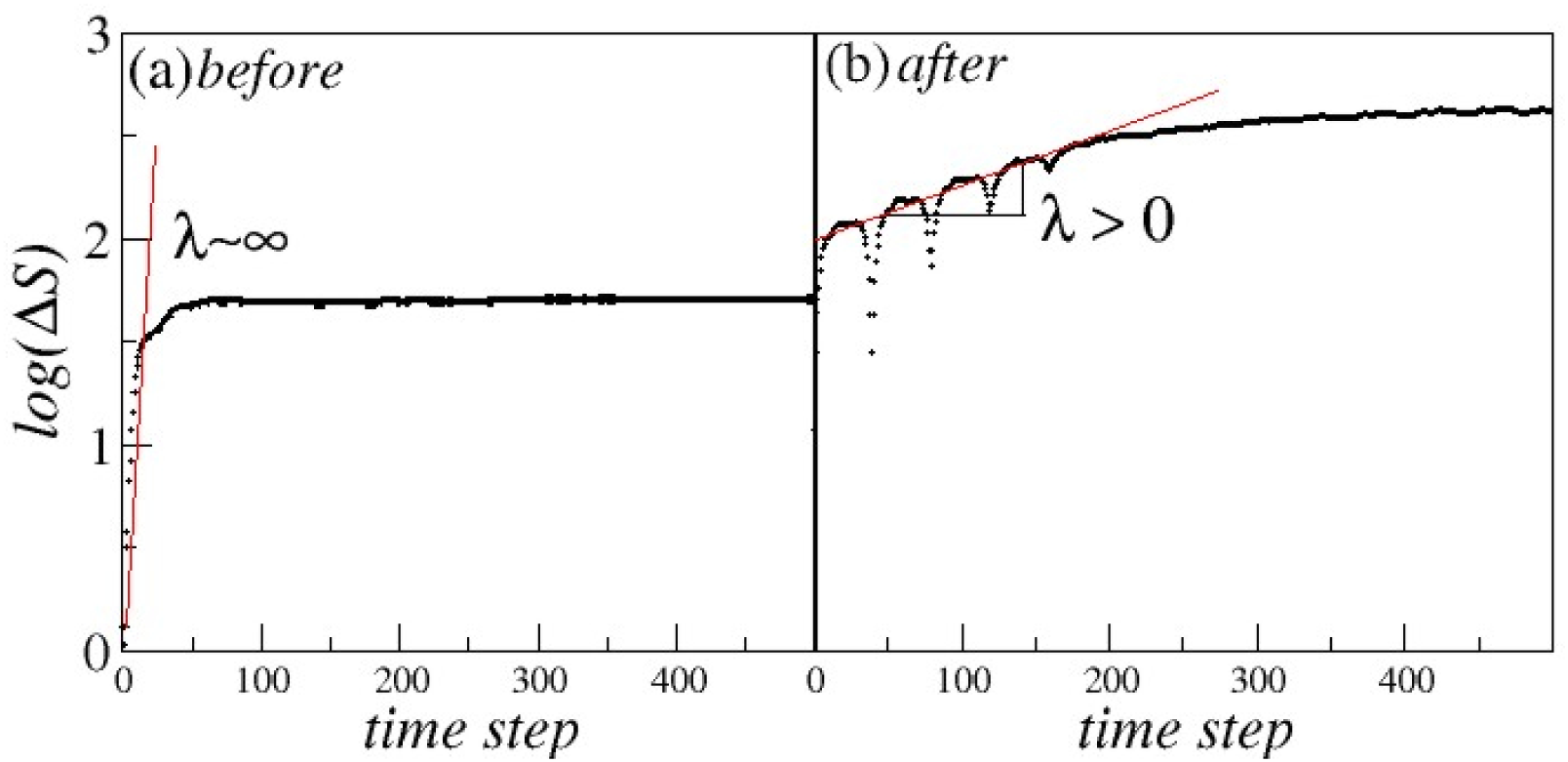
Logarithm of the divergence due to the time step for the same plant time series before and after the stimulus. In (a) it was observed an approximately infinite exponent showing that the series was stochastic. In (b), for the same plant after stimulus, the exponent was finite and positive, indicating that the dynamics has become chaotic.

Further, analysing visually and systematically the shape of each original time series before and after stimuli, we have observed the presence of spikes up to 500 μV (taking into account the */ΔV/* baseline range around 20 – 40 μV) in all time series after stimuli (inserts in Figure 6). Then, the distribution of the */ΔV/* magnitudes was analysed in the whole scored time series after stimuli (aprox. 1 h of sampling), in order to obtains a more accurate and reliable analysis. Surprisingly, but not unlikely, a power law distribution was detected, *D(/ΔV/)* = */ΔV/^-μ^*, with exponent μ ≅ 2.0 (Figure 6). This result indicates that the time series exhibits scale invariance^19^.

**Figure 6.**
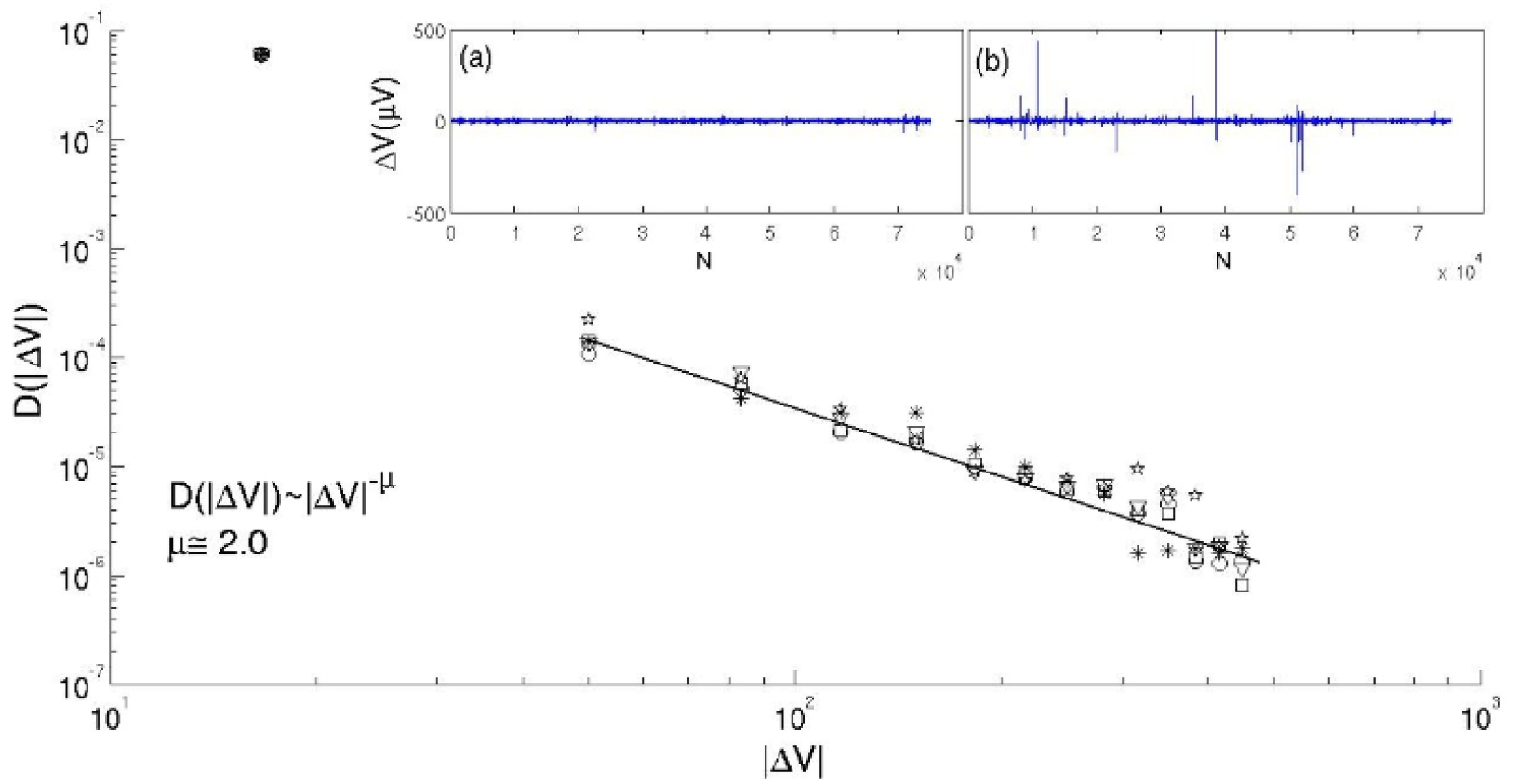
Distribution of */ΔV/* μV in the asymptotic limit from five time series after stimuli following a power law, *D(/ΔV/)*=*/ΔV/^-μ^*, whit μ values from 1.9 up to 2.1. Representative samples from the original time series scored before and after stimuli are showed in the insert (a) and (b), respectively.

## Discussion

Plants, as any live being, are thermodynamical dissipative open systems far from equilibrium, with self-organized emergent properties^36,37^. Such systems often show non-linear dynamics that can fit to deterministic chaos^18,19^. Chaotic dynamics have been observed in many biological phenomena^38,39^, ranging from human EEG^18^, different processes in plant physiology^24,40,41^, to ecological dynamics^42^. Our results consistently showed that the temporal dynamics of electrical signaling in plants is highly complex (but not random), indeed chaotic. As far we can know, this is the first observation of chaos in plant electrical signaling.

Thus, some questions arise: What could engender such complexity? Is there some physiological meaning in such chaotic behavior? Although plants do not have a nervous system, they possess a complex network that uses ion fluxes moving through defined cell types to rapidly transmit information between distant sites within the organism^43^. Distinction between APs and VPs in plants based solely upon their kinetic characteristics might be doubtful because VPs often mimic AP kinetics, and they often overlap each other^14,44^, arising a spatial and temporal complex web of electrical signaling^4^. Additionally, the well known complexity of calcium waves underlying plant electrical signaling^13,42^, and the network of neurotransmitters signaling, acting on development, communication and stress responses^5,17^ can explain, at least in part, the temporal complexity in the electrical transmission of significant (not random) physiological information through the plant body, which is not properly congregated by single VPs or APs signals^14^.

Furthermore, several electrical signals from single cells can be synchronized in time and space^4,15^, similarly to a neuronal electrical network. Our finding that the distribution of *ΔVs* follows a power law can bring some light to the phenomenon of synchronization. Power law distributions, specifically when 1 < μ < 3 (in our results μ was between 1.9 and 2.1), can be signatures of self-organized critical (SOC) systems^45–47^. SOC is a ubiquitous phenomenon in nature regardless on the details of the physical system under study observed, for instance, in earthquakes, sandpiles, droplet formation, dynamic of populations, and in biological evolution^46,47^. Thus, we hypothesized that the spikes observed herein (Figure 6) and those reported by Masi et al. (ref 15) emerge from a self-organized process, when groups of cells with different sizes synchronize their variation of voltages. Moreover, because the exponent μ is around 2.0, the distribution of spikes has not a characteristic size^20^.

According to Bak et al. (ref 45), SOC systems are barely stable and, then, perturbations can cause a cascade of energy dissipation on all length scales, following a power law. Stability is supposedly linked in a straightforward way with the system dynamics. In a biological interpretation, it is expected that more complex dynamics allow system stability, providing higher resilience to the system under external disturbances^25,48^. Thus, the higher the complexity of the system is, “healthier” it shall be. For instance, plants showing more complex dynamics in leaf gas exchanges under control conditions showed better recovering after a water deficit situation^24,25^. Studies with EEG in human brains have also found a correlation between complexity and health. For example, dramatic reductions in the complexity of EEG signals have been associated with seizures in humans^22^. Similarly, analysis of the temporal dynamics of EEG in Alzheimer patients identified a reduction in irregularity (complexity) of the time series of electric signals^20^. Accordingly, our results from complexity measurements after stimuli indicated that the osmotic stimulus was stressful, reducing the complexity and stability, pushing the system to a critical state.

Recent studies with EEG in human brains have also showed evidences of SOC^49,50^. However, SOC has been associated to a normal brain functioning and disturbances, such as epileptic seizures attacks, deviate neuronal activity from a power-law distribution^50^. Thus, while SOC seems to be the normal state of brain functioning, our results suggest that the critical state in plants can be reached under stressful environmental conditions. Therefore, it is not clear what is the meaning of SOC for biological systems stability. By one hand, critical states can be associated to optimal information processing and computational capabilities (brain specialties)^50^ and, on the other hand, in the critical state, systems can dissipate energy (tensions) efficiently^45^, which is a very important capacity for plants under stressful situations.

### Follow-up questions and perspectives

Summarizing, our results support the hypothesis that, beyond APs and VPs, there is a complex and significant, i.e. non-random and responsive to stimuli, flow of electrical information, showing chaotic dynamic. Additionally, when disturbed, the system can reach a self-organized critical state.

Our results suggest that electrical signaling could play a broader role in plant life. The complexity in the electrical signals measured here supports a further hypothesis that a large amount of meaning information can flow throughout the plant body, likely affecting the signaling of a plethora of processes. Moreover, since the complexity of the electrical signals can be affected by environmental stimuli, it is reasonable to assume a special involvement in plant stress responses, such as single APs and VPs supposedly do^6,8^.

As these are the first reports of chaos and SOC phenomena in plant electrophysiology, we should ask if the underlying electrical information could be directly related to the plant responses to specific environmental stimuli, and if different cultivars/species could show different response patterns. Additionally, we shall ask: Which are the causal mechanisms of SOC in the electrical signals? What could be the physiological significance of critical state for the plants? In order to answer these fundamental questions, more experimental studies are necessary to support the development of mathematical models, allowing explore deeply the origins of SOC in plant electrophysiology.

Further, if there is ubiquitous complex information underlying the electrical signals in plants, we could decode this information and use it to create a protocol of diagnosis, gathering their physiological states in real time. Practically, we could try to uncover a plant’s “language”.

## Acknowledgments

This work was supported by Stoller do Brasil. GMS is research fellow of the CNPq (Conselho Nacional de Pesquisa e Desenvolvimento Tecnológico); GFRS is PhD fellow of the CAPES (Coordenação de Apoio de Pessoal do Ensino Superior). We wish to thank Prof. Alexander G. Volkov for suggestions and commentaries on the manuscript, and to Prof. Luiz Gonzaga E. Vieira for the text review.

## Author contributions

G.F.R.S. designed and performed research; A.S.F. analyzed data; G.M.S designed research and wrote the paper. All authors discussed and interpreted the results.

